# RAmpSim: A Thermodynamic Simulator for Hybridization Capture in Metagenomic Sequencing

**DOI:** 10.64898/2025.12.05.692407

**Authors:** Aidan Zhang, Christina Boucher, Noelle Noyes, Yun William Yu

## Abstract

Hybridization (bait) capture combined with long-read sequencing enables targeted profiling within complex metagenomes but introduces systematic biases from bait multiplicity, sequence composition, and species abundance that existing simulators ignore. We present `RAmpSim`, a fast simulator that models bait–target hybridization and fragment capture using a thermodynamic nearest-neighbor energy model and Boltzmann-weighted sampling of binding sites. Fragments are generated through multinomial sampling parameterized by bait concentration, binding energy, and genomic abundance before being passed to existing long-read simulators for modeling platform-specific errors. Implemented in Rust, `RAmpSim` reproduces empirical within-genome coverage and cross-species enrichment patterns observed in capture-based metagenomic datasets. Compared to uniform-coverage baselines, `RAmpSim`’s simulated coverage distributions are up to an order of magnitude closer to real data with respect to earth mover’s distance. Classification analysis reveals high recall in classifying high coverage regions between simulated and experimental distributions while outperforming a uniform baseline. Supporting accurate benchmarking and bait-set evaluation, `RAmpSim` provides an interpretable, efficient framework for simulating capture-based metagenomic sequencing.

**Code Availability:** https://github.com/az002/RAmpSim.git

## 1 Introduction

Shotgun metagenomics has significantly advanced our understanding of microbial communities and their interactions with hosts in a variety of environments and samples, including soil and water in agricultural environments, clinical specimens from human patients, and wastewater used for population-scale surveillance [15]. One of the ongoing challenges that arises with shotgun metagenomics is that genes and genomic regions of interest are frequently dwarfed by extraneous DNA from other microbial species (such as bacteria) or hosts (such as humans). For example, Noyes et al. [10] demonstrated that antimicrobial resistance genes (ARGs)—which define the resistome of a sample—can comprise less than 1% of the sequence data within a shotgun metagenomic sample. This greatly limits our ability to resolve the diversity and composition of such samples, and in practice often necessitates substantially higher sequencing depth or targeted enrichment (e.g., host depletion or capture-based approaches) to recover low-abundance targets with confidence.

Bait-capture, also known as hybridization capture and target enrichment, is a laboratory method for selectively enriching nucleic acid fragments prior to sequencing. In a typical workflow, a complex DNA or cDNA library is incubated with a pool of oligonucleotide baits that are designed to hybridize to regions of interest; bait-target hybrids are then isolated, washed to remove non-hybridized molecules, and prepared to produce a sequencing library that is enriched for the targets. See Bravo et al. [1] for a more detailed discussion of the laboratory procedure. This approach increases sensitivity for low-abundance genes or genomic regions, such as ARGs as discussed above. By limiting off-target reads, sequencing costs are also greatly reduced, enabling more focused studies. For example, bait-capture has been used to study many types of genomic regions, including specific exomes, bacterial species, and viruses, to name a few. Recently, in target-enriched long-read sequencing (TELSeq), bait-capture is combined with long-read sequencing, which further improve the detection sensitivity of rare genes [17].

A central challenge in studying bait-capture and developing robust computational methods for the resulting data is the scarcity of ground truth. True target presence, exact on-target fractions, and bias profiles are almost never known for real samples. Simulation provides a practical remedy to this problem through the synthetic generation of a ground truth that can be used to estimate precision and recall. Hence, by generating synthetic datasets with known compositions, various bait designs and novel methods can be systematically evaluated. This use of synthetic data has been paramount to progress in bioinformatics. Simulated reads have routinely enabled the benchmarking of genome assemblers, structural variant callers, and RNA-seq transcript quantification methods [21, 13, 12]. Given this importance, there has been a plethora of work in developing accurate and robust read simulators. ART and pIRS were some of the earliest read simulators to simulate Illumina reads from whole genome sequencing experiments [4, 3]. BEAR was one of the earliest shotgun metagenomics read simulators [5]. PBSIM3 and NanoSim simulate the read length and error profiles of Pacific Biosciences and Oxford Nanopore long-read sequencing data [11, 19]. In these settings, simulators often assume that reads are sampled uniformly from reference genomes, which typically holds true for untargeted sequencing protocols, including shotgun metagenomics. Although it is possible to design software that works on enriched reads [16], such software is typically benchmarked on less complexly simulated datasets; their performance on bait-capture enriched reads has in the past only been able to be assessed qualitatively.

In bait-capture protocols, the express purpose is to induce a non-random sampling of genomic loci, thus breaking the uniform-sampling assumption. Bait-target hybridization, washing, and amplification create predictable but uneven coverage patterns. The coverage of the reads usually peaks where the baits tile the genome (i.e., in the targeted regions) and falls off toward the bait edges (i.e., in the non-targeted regions). Enrichment efficiency can also vary substantially with the GC content, fragment length, and other laboratory conditions (e.g., temperature). In addition, near-homologous sequences can lead to off-target fragments being captured and sequenced. In metagenomic samples, these effects compound with abundant host DNA, wide species abundance ranges, and species-specific bait affinities. This bias creates a substantial mismatch between the assumptions of the existing read simulators and the characteristics of the real data. Consequently, benchmarking tools and algorithms on synthetic reads generated under uniform-coverage assumptions from these simulators may not reflect their performance in targeted capture datasets. Methods such as PBSIM3 and NanoSim are capable of producing non-uniform coverage patterns through empirical sampling, however these data-driven approaches require substantial training data, generalize poorly to other laboratory protocols, and offer limited interpretability [11, 20].

In this paper, we introduce a read simulator that models the thermodynamic interactions between baits and target fragments. We refer to our method as `RAmpSim` (Read Amplification Simulator). By explicitly incorporating relative bait concentrations, binding energy, and genomic abundance, `RAmpSim` generates fragment distributions that seamlessly integrate with existing read simulators to produce realistic bait-capture data. Using publicly available datasets, we demonstrate that our method reproduces realistic coverage patterns within and between species in a mock community sample. The resulting simulator provides a reproducible and interpretable foundation for benchmarking bait-capture metagenomic workflows and optimizing bait designs.

## 2 Methods

### 2.1 Simulation Overview

Assume that we have a set of baits with with known relative concentrations in solution, a set of reference genomes, and additional hyperparameters determining hybridization energetics and the distribution of fragment lengths. We outline the computational steps for simulating bait-based hybridization capture prior to sequencing through the following stages:

#### 1. Bait-Target Matching

Baits are mapped to each reference genome using sequence alignment. This gives a set of possible hybridization sites across reference genomes.

#### 2. Energy-Based Scoring

Each hybridization site is scored with its binding free energy (*ΔG*) estimated using a Thermodynamic Nearest-Neighbors model (TNN) with SantaLucia parameters [14]. For simplicity, mismatches in our model are set to have *ΔG* = 0.

#### 3. Fragment Sampling

Through multinomial sampling steps, capture fragment centers are drawn based on the relative concentration of baits near it, their binding probabilities, and the abundance of the reference. Background fragments are sampled uniformly within genomes with coverage depending on genomic abundance. All fragment lengths are determined following a user-defined log-normal distribution.

#### 4. Error Modeling

Sequencing errors are introduced using existing read simulators such as PBSIM3 with platform-specific models designed to replicate empirical error profiles for generating realistic reads [11].

#### Hybridization Model

Consider a metagenomic sample with reference genomes 𝒢 and abundances {*a*_*g*_} _*g*∈𝒢_ . Let ℬ ⊂ *∑*^∗^ be the set of oligonucleotide baits over the nucleotide alphabet *∑* = {*A, C, G, T*} . For baits *p* ∈ ℬ, let *H*_*p*_ be the set of candidate hybridization sites (*x, g*) where *x* is the position-indexed sequence of the site within its corresponding reference genome *g* ∈ 𝒢.

Given a thermodynamic model *E*(*x, p*) that estimates *ΔG* of hybridization between *x* and *p*, we define the unnormalized binding score:

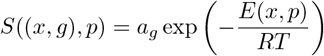

where the exponential term is the Boltzmann factor, which is an unnormalized relative probability describing the propensity of bait-target binding at equilibrium. In particular, it is a function of the energy *E*(*x, p*) and the temperature of the system *T* during hybridization, where *R* is the universal gas constant relating temperature to energy. The abundance term *a*_*g*_ weights the score according to the relative representation of the reference in the sample.

In the TELSeq protocol, PCR amplification is performed prior to hybridization, resulting in high fragment multiplicities relative to bait concentrations. In this setting, bait-bait competition can be ignored and bait hybridization can be viewed as saturated. We model the competition among the binding sites for a fixed bait *p* by normalizing over all candidate sites:

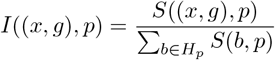

Here *I*((*x, g*), *p*) represents the expected fraction of baits *p* bound to the site (*x, g*). This is analogous to the formula for fractional occupancy under the Langmuir isotherm model, which is commonly used in DNA microarray hybridization models [7, 2]. However, in this case, the roles of baits and target fragments are reversed: fragments are viewed as fixed, whereas baits are free to bind to sites distributed among the fragments.

#### Fragment Sampling

We can model the generation of fragments by drawing from a series of multinomial distributions. Since we assume that bait-bait competition is negligible, we model the overall binding rate of each bait *p* as proportional to its relative concentration *c*_*p*_ with a shared rate constant. A Poisson model fitted on these rates is used to model the counts of binding events for each bait.

Treating a bait binding event as the capture of a fragment, if we condition on the number of fragments *N*, then the overall process is equivalent to sampling from a multinomial

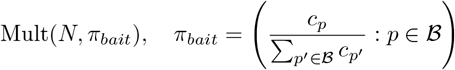

Similarly, for a bait *p*, we model the rate at which it binds to candidate sites *b* ∈ *H*_*p*_ as proportional to the score *S*(*b, p*), which parameterizes a Poisson distribution. Thus, after conditioning on the number of binding events with bait *p* as *N*_*p*_ = *Nw*_*p*_, we obtain another multinomial

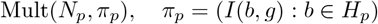

After drawing the binding site (*x, g*), we draw a fragment length under an empirically determined user-defined read length distribution, *L* ∼ Lognormal(*µ, σ*^2^). This determines a uniform distribution of fragment centers in a window of length *L* centered on the binding site from which the actual fragment is ultimately drawn.

To sample background fragments for a fixed number of fragments *N* ^*′*^, we sample *a*_*g*_*N* ^*′*^ fragments for each genome *g* ∈ 𝒢 uniformly at random across all positions in the genome.

### 2.2 Implementation Details

`RAmpSim` is implemented in Rust for computational efficiency. All computations are performed on a single CPU thread. Reference genomes are loaded into memory for fast access during bait–target evaluation and fragment sampling.

As the primary input, `RAmpSim` accepts a SAM file containing candidate bait–target alignments generated from read mappers such as Bowtie2 [6]. Our mapping criteria excludes indels and allows for at most 40 mismatches. From these alignments, fragments are sampled according to the hybridization and fragment-length models. `RAmpSim` has parameters that determine the fraction of capture compared to the background fragments, the genomic abundances of the present species, the hybridization temperature (used in the Boltzmann-weighted energy model) and the log-normal fragment-length distribution ln *µ* and ln *σ*. These parameters can be given by the user or estimated from sequencing libraries.

Both capture-derived and background fragments are outputted as a FASTA file. On TELSeq-scale datasets ( ∼10000 baits producing ∼100000 fragments), a complete simulation runs in a few seconds on a workstation equipped with an Intel Xeon Gold 5416S CPU and 512 GB of RAM; however, in our simulations of microbial genomes, the peak memory usage was less than 100 MB.

### 2.3 Simulation Setup (Hyperparameter Tuning)

To validate `RAmpSim`, we used publicly available TELSeq datasets sequenced from a vendor supplied mock microbial community (ZymoBIOMICS Microbial Community DNA Standard II [Log Distribution]) including a mixture of bacterial and fungal species that decrease in genomic abundance by a factor of 10 (Table 1) [17]. The theoretical abundances are used to weight the bait binding scores described in the hybridization model and determine the relative proportions of background fragments generated from each species.

**Table 1.** Composition of ZymoBIOMICS Microbial Community Standard II. The genomic abundances for each species is used as the weighting term *a*_*g*_ for the binding scores in the hybridization model. The abundances also directly determine the relative proportions of sampled background fragments.

| Organism | Genomic Abundance (%) |
| --- | --- |
| *Listeria monocytogenes* | 89.100000 |
| *Pseudomonas aeruginosa* | 8.900000 |
| *Bacillus subtilis* | 0.890000 |
| *Saccharomyces cerevisiae* | 0.890000 |
| *Escherichia coli* | 0.089000 |
| *Salmonella enterica* | 0.089000 |
| *Lactobacillus fermentum* | 0.008900 |
| *Enterococcus faecalis* | 0.000890 |
| *Cryptococcus neoformans* | 0.000890 |
| *Staphylococcus aureus* | 0.000089 |

The TELSeq dataset includes reads from three mock community samples, each independently physically fragmented to achieve different target median fragment length distributions (2kb, 5kb, and 8kb) prior to sequencing. To focus on modeling fragment capture, we remove the effects of PCR bias by deduplicating reads using samtools markdup [8].To fit the distribution of fragment lengths, the scipy stats package was used to determine the *µ* and *σ* parameters of a log-normal distribution for each dataset [18]. To estimate the proportion of capture-derived and background fragments, reads within *µ/*2 of candidate bait binding sites were considered capture reads and the rest as background, according to the length *L* window used to generate fragments in `RAmpSim`. Lastly, the temperature for each dataset was set to 70 ^*°*^C based on the protocol outlined by Slizovskiy et al. [17]. Histograms of fragment length distributions with fitted log-normal curves and a table of capture to background fragment ratios with other model hyperparameters are shown in Supplementary Figure S1 and Supplementary Table S1 respectively.

The generated fragments are inputted into PBSIM3 using the template strategy and ERRHMM-Sequel model. We simulate reads with one pass per molecule, which yields a higher error rate than the≈ 5-pass CCS reads in the TELSeq datasets. Our evaluation focuses on metrics based on fragment locations and multiplicities such as coverage EMD, peak recall, and genomic abundance. We do not expect this discrepancy in error rate to materially affect our conclusions as these metrics are robust against per-base accuracy.

## 3 Results

### 3.1 Fidelity of Coverage Patterns

The simulated coverage profiles from our method closely reproduce the empirical normalized coverage patterns for large contiguous genomes across all replicates at log scale. For both simulated and experimental data, coverage levels are highest near regions with high bait-density and decays to background levels otherwise (Figure 2). Coverage distributions for additional genomes and replicates are shown in Supplementary Figures S2 and S3.

**Fig. 1.**
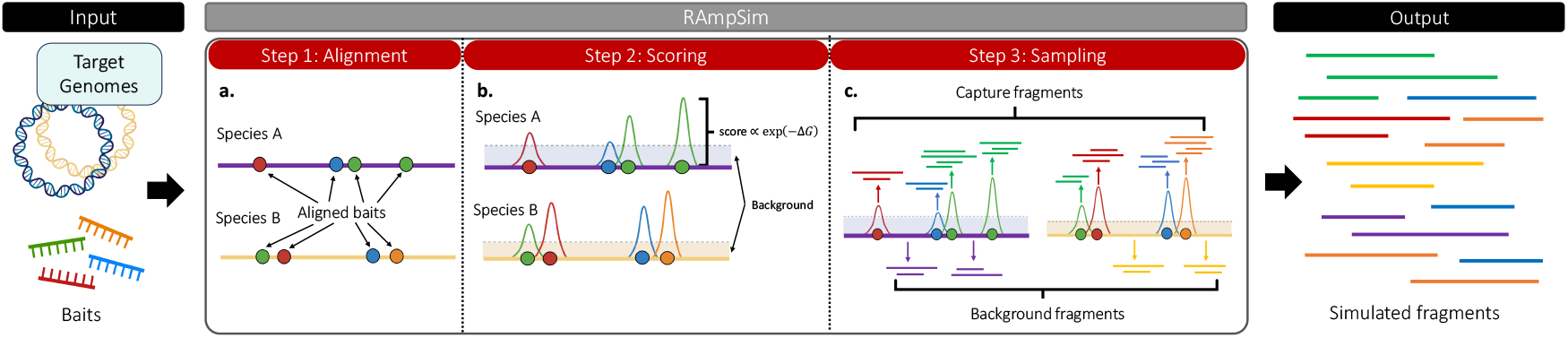
Simulation framework for generating fragments. **a**. Bait sequences are aligned to each reference genome present within the sample, allowing secondary mappings to capture off-target hybridization sites. **b**. Candidate binding sites discovered from the alignment step are scored using its hybridization free energy (*ΔG*). Energies across candidate sites for a given bait are normalized with the sum of their abundance weighted scores. **c**. Capture fragments are sampled according to multinomial distributions on baits and their binding sites while background fragments are sampled uniformly across positions.

**Fig. 2.**
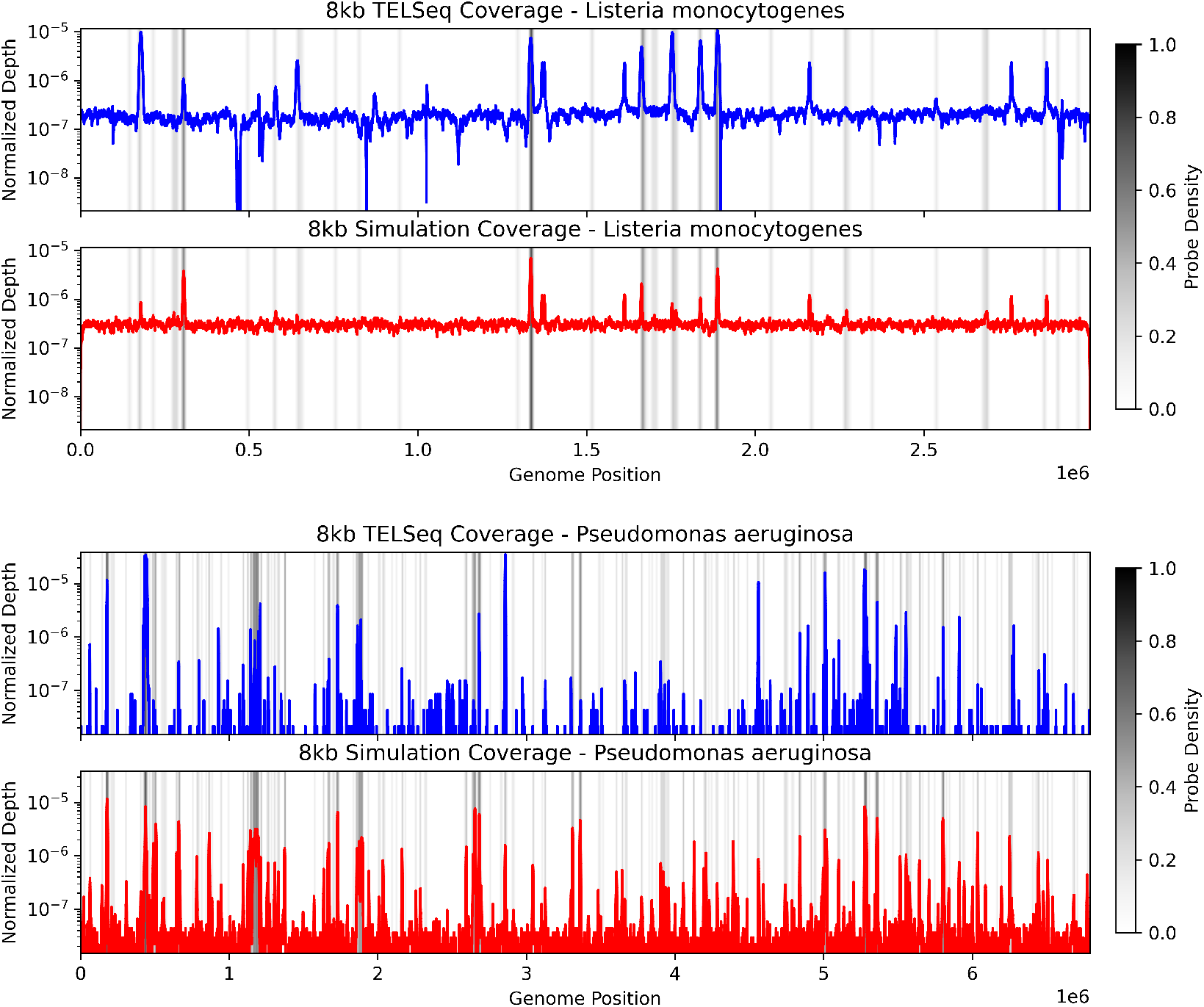
Log coverage distribution plots for *L. monocytogenes* (top) and *P. aeruginosa* (bottom). The coverage distribution for each genome was normalized by the sum of the coverage over each position. The coverage patterns between the observed (blue) and simulated (red) data are similar, in which peaks in each distribution correspond to high bait-density regions shown in gray. Outside of these regions, coverage falls down to background levels. Coverage distributions for additional genomes and replicates are shown in Supplementary Figures S2 and S3.

We quantify the similarity between the observed and simulated coverage distribution using the earth mover’s distance (EMD), which can be viewed as the minimum cost to transform one distribution to the other, where the cost is proportional to the amount of density weighted by the distance it is moved. The EMD between the empirical and simulated coverage distributions is consistently lower than that between the empirical coverage distribution and a uniform baseline, but the difference is smallest for *L. monocytogenes* and *L. fermentum* (Table 2). However, this is expected since the observed coverage distribution of *L. monocytogenes* is mostly uniform with sparse peaks, which the simulated data reproduces. For *L. fermentum*, we note that the original bait designs did not target any region in its genome due to its lack of ARGs so that its coverage distribution is dominated by background reads, which would also result in a uniform coverage distribution. EMD results for additional replicates are provided in Supplementary Table S2.

**Table 2.** EMD results for the 8kb replicate. For prokaryotic genomes with single chromosomes, the EMD between the empirical and simulated coverage distributions outperforms the baseline. Eukaryotic species such as *S. cerevisiae* and *C. neoformans* contain multiple chromosomes of varying lengths and were not considered targets in the original bait design. Tables for additional replicates are shown in Supplementary Table S2.

| Species | EMD(tels, sims) | EMD(tels, unif) |
| --- | --- | --- |
| *Listeria monocytogenes* | **2.01e-07** | 2.70e-07 |
| *Pseudomonas aeruginosa* | **1.45e-07** | 2.74e-07 |
| *Bacillus subtilis* | **9.75e-08** | 4.51e-07 |
| *Escherichia coli* | **1.76e-08** | 3.21e-07 |
| *Enterococcus faecalis* | **3.37e-07** | 6.78e-07 |
| *Lactobacillus fermentum* | **7.30e-07** | 1.05e-06 |
| *Salmonella enterica* | **6.91e-08** | 3.46e-07 |
| *Staphylococcus aureus* | **4.47e-07** | 7.28e-07 |

To evaluate the simulator’s ability to recover the locations of high coverage, the problem can be framed as a binary classification task. Specifically, positions within a genome are classified as high coverage based on whether the raw coverage at the position passes a median threshold computed from all positions in the genome. This procedure is performed on both the observed and simulated data, yielding separate labeled sets of locations for comparison with the observed high coverage locations being considered ground truth. For most genomes, the F1 score is at least 0.5 with the precision being consistently higher than the recall (Table 3). The high precision shows that `RAmpSim` produces high coverage regions that coincide with those in the observed data. The lower recall suggests that many positions classified as high coverage in the TELSeq data fall below the median threshold in the simulated coverage, especially for low-abundance species. In particular, this indicates that `RAmpSim` only recovers a subset of the high coverage positions observed experimentally.

**Table 3.** F1, precision, and recall scores of high/low coverage classification by `RAmpSim` and a uniform baseline in the 8kb replicate. Uniform baseline values are reported as mean *±*95% confidence intervals over *N* = 100 simulated uniform-coverage datasets. For each species, the left block reports the scores obtained by `RAmpSim`, and the right block reports the corresponding uniform-baseline metrics. For all species except *L. monocytogenes*, the F1, precision, and recall for the high coverage regions called by `RAmpSim` lie above the 95% confidence intervals of the uniform baseline. See Supplementary Table S3 for corresponding F1 metrics on the 2kb and 5kb replicates.

| Species | RAmpSim |  |  | Uniform baseline |  |  |
| --- | --- | --- | --- | --- | --- | --- |
|  | F1 Score | Precision | Recall | F1 Score | Precision | Recall |
| *Listeria monocytogenes* | **0.5030** | **0.5119** | 0.4944 | 0.477 ± 0.024 | 0.480 ± 0.030 | 0.474 ± 0.024 |
| *Pseudomonas aeruginosa* | **0.3152** | **0.5312** | **0.2240** | 0.233 ± 0.019 | 0.379 ± 0.031 | 0.169 ± 0.014 |
| *Bacillus subtilis* | **0.3678** | **0.5524** | **0.2756** | 0.287 ± 0.027 | 0.487 ± 0.052 | 0.204 ± 0.019 |
| *Escherichia coli* | **0.7661** | **0.7879** | **0.7455** | 0.374 ± 0.027 | 0.402 ± 0.030 | 0.350 ± 0.024 |
| *Salmonella enterica* | **0.7202** | **0.8518** | **0.6239** | 0.347 ± 0.023 | 0.480 ± 0.032 | 0.271 ± 0.018 |
| *Lactobacillus fermentum* | **0.0351** | **0.4105** | **0.0183** | 0.000 ± 0.000 | 0.000 ± 0.000 | 0.000 ± 0.000 |
| *Enterococcus faecalis* | **0.4995** | **0.9813** | **0.3350** | 0.058 ± 0.041 | 0.189 ± 0.133 | 0.034 ± 0.024 |
| *Staphylococcus aureus* | **0.1743** | **0.9878** | **0.0956** | 0.033 ± 0.053 | 0.034 ± 0.055 | 0.033 ± 0.048 |

We also compare the F1, precision, and recall against a random uniform baseline. For *N* = 100 iterations, random uniform distributions were generated by sampling fragment start positions uniformly across each genome, with fragment lengths drawn from the fitted lognormal distributions. The number of fragments was chosen so that the expected mean coverage matched the empirical average coverage depth. We then applied the same coverage-classification pipeline to these simulated coverages and summarized the resulting metrics as mean *±*95% confidence intervals. For most species, the F1, precision, and recall achieved by `RAmpSim` are higher than the corresponding uniform-baseline intervals, whereas performance for *L. monocytogenes* is only modestly better than the baseline, consistent with its coverage distribution being close to uniform. Additional F1 metrics for the 2kb and 5kb replicates are reported in Supplementary Table S3.

To make differences between the simulated and observed coverage distributions more apparent, we plot the coverages on a common scale, with discrepancies especially pronounced for *P. aeruginosa*. In particular, prominent coverage peaks at approximately 0.5 and 3 Mb in the experimental data are markedly attenuated in the simulated data (Figure 3). This may be partially explained by the prominence of the background distribution pulling away density from peaks in the simulated coverage profile.

**Fig. 3.**
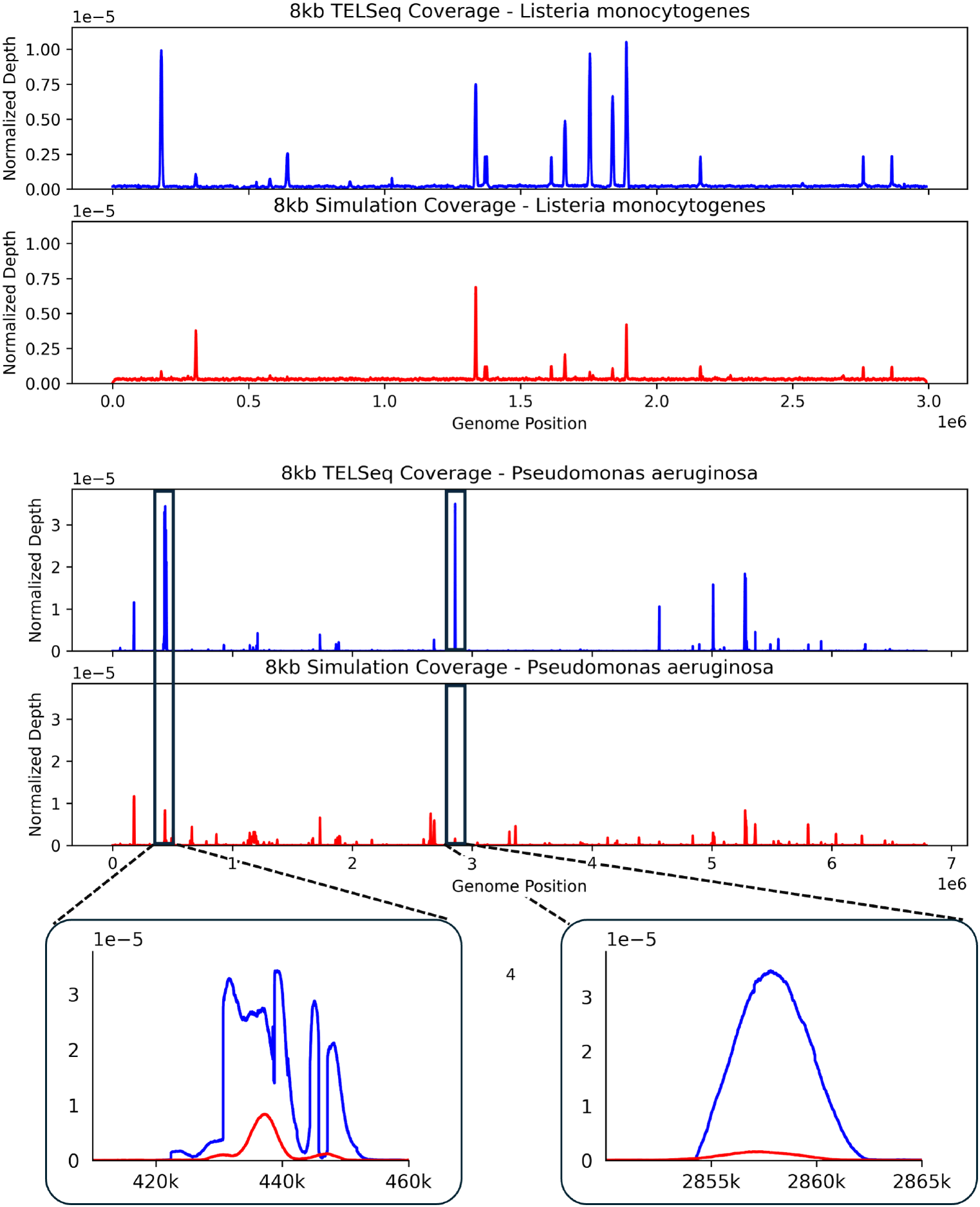
Normalized coverage distributions for *L. monocytogenes* (top) and *P. aeruginosa* (bottom). Although the simulator accurately reproduces the locations of peaks, the relative heights of peaks can differ substantially between the simulated and experimental data. We show two locations in the *P. aeruginosa* genome ( ∼ 440 kb and ∼ 2.86 Mb) where the difference is largest. Coverage distributions for additional genomes are shown in Supplementary Figures S2 and S3.

### 3.2 Fidelity of Species Abundances

In addition to within-species coverage patterns, we evaluate `RAmpSim`‘s ability to reproduce the abundance of species observed in the observed data relative to the theoretical genomic abundances. Abundances were computed by assigning reads to species based on unique QC-filtered minimap2 alignments outlined by Nicholls et al. [9].

In general, the simulated and observed data show similar abundance patterns (Table 4). For example, in both datasets, most reads map to *L. monocytogenes*, which has the highest theoretical genomic abundance in the mock sample. Furthermore, target species such as *E. coli, S. enterica*, and *B. subtilis* in both the simulated and observed data show substantially increased abundance compared to the theoretical compositions as the bait design for the community selectively targets ARGs present in each genome; non-target species (due to their lack of ARGs) such as *S. cerevisiae, C. neoformans*, and *L. fermentum* either showed suppressed abundance or remained having the lowest abundance relative to other species. Additional tables for the remaining replicates are shown in Supplementary Table S4.

**Table 4.** Comparison of theoretical, observed, and simulated abundances of the 8kb replicate. In general the changes in genomic abundances compared to the theoretical values are preserved between the observed and simulated data. *L. monocytogenes* remains the most abundant species, and species targeted by the bait design such as *B. subtilis, E. coli*, and *S. enterica* are enriched; non-target species, including *L. fermentum, C. neoformans*, and *S. cerevisiae* are depressed or remain having the lowest abundance. The main discrepancies are the absolute changes between the simulated and observed abundances, being most prominent for *L. monocytogenes* and *P. aeruginosa*. See Supplementary Table S4 for the abundance comparisons of remaining replicates.

| Species | Theoretical | Observed | Simulated |
| --- | --- | --- | --- |
| *Listeria monocytogenes* | 89.100000 | 64.893325 | 35.984226 |
| *Pseudomonas aeruginosa* | 8.900000 | 9.759034 | 26.771994 |
| *Bacillus subtilis* | 0.890000 | 9.183611 | 2.717070 |
| *Saccharomyces cerevisiae* | 0.890000 | 0.090746 | 0.370099 |
| *Escherichia coli* | 0.089000 | 10.069688 | 14.768590 |
| *Salmonella enterica* | 0.089000 | 5.964689 | 15.031289 |
| *Lactobacillus fermentum* | 0.008900 | 0.000993 | 0.189728 |
| *Enterococcus faecalis* | 0.000890 | 0.026362 | 1.739722 |
| *Cryptococcus neoformans* | 0.000890 | 0.000000 | 0.000000 |
| *Staphylococcus aureus* | 0.000089 | 0.010560 | 2.237555 |

Although the overall abundance patterns between the simulated and observed data are similar, there are nontrivial differences in the absolute abundance values of species between the datasets. For example, the abundance of *L. monocytogenes* is much lower in the simulated data compared to the observed data, whereas species such as *P. aeruginosa* and *S. enterica* are even more enriched in the simulated data. In particular, *P. aeruginosa* is known to have high GC content leading to inefficiencies in DNA extraction, which could explain the substantial increase in abundance in the simulated data compared to the observed data.

## 4 Discussion

The results show that `RAmpSim` is able to recover large-scale coverage structures observed in bait-capture sequencing experiments. For the majority of bacterial genomes with a single main chromosome, simulated coverage peaks coincide with high bait-density regions and decay to background elsewhere, matching the empirical profiles reported in Figure 2 and Supplementary Figures S2 and S3. Together with the low earth mover’s distances between simulated and observed coverage distributions (Table 2; Supplementary Table S2), this suggests that a model that combines alignment-derived candidate hybridization sites, thermodynamics-based hybridization scores, and abundance-weighted sampling is sufficient to explain the dominant enrichment patterns in this dataset.

The main discrepancies between simulated and experimental data arise in two settings: (1) peaks present in the experimental data but absent or attenuated in the simulated data, and (2) differences in the relative heights of peaks within the same genome (Figure 3; Supplementary Figures S2 and S3). Both effects reflect a smoothing of coverage distributions generated from `RAmpSim` relative to observed data. In particular, high coverage peaks in the TELSeq coverage appear as more moderate peaks in the `RAmpSim` coverage, while the background coverage in regions between peaks that are more sparse in the TELSeq data appear with higher coverage in the `RAmpSim` data. In practice, these differences can be attributed to model components that were intentionally simplified. GC-dependent effects on hybridization and amplification were not explicitly modeled, even though high-GC regions are known to be more difficult to denature and capture. A broader background component in the simulator also increases the baseline signal across positions, which can flatten the normalized coverage distributions and reduce contrast among peaks.

Coverage classification analysis results are consistent with this view. Precision values are high across most genomes, indicating that high coverage locations called by the simulator correspond to true high-coverage regions (Table 3; Supplementary Table S3). Recall is lower, especially for low-abundance or non-target species such as *L. fermentum*, where any observed coverage is sparse and often below the median-based threshold. This pattern is expected when the underlying experimental data contain weak off-target signals that are incompletely captured by the alignment-derived candidate sites or are down-weighted by the hybridization model. Overall, the dominant failure mode of `RAmpSim` is estimating the intensity of coverage, rather than systematically misplacing coverage.

To assess whether this performance could be explained by random fragmentation alone, we compared F1, precision, and recall to a uniform baseline in which fragments were sampled uniformly along each genome, with lengths drawn from the fitted fragment-length distributions and the number of fragments chosen to match the empirical mean coverage. Across most bacterial species, the F1, precision, and recall of `RAmpSim` lie above the corresponding 95% confidence intervals of the uniform baseline (Table 3; Supplementary Table S3), indicating that the observed agreement with TELSeq high coverage positions cannot be attributed solely to random noise under uniform coverage. For low-abundance species such as *E. faecalis* and *S. aureus*, the uniform baseline yields very low F1 scores, whereas `RAmpSim` still recovers a subset of experimentally observed high coverage locations with high precision, consistent with the simulator producing genuine positional signal in the TELSeq profiles. In contrast, performance for *L. monocytogenes* is only modestly better than the uniform baseline, in agreement with its small EMD and the fact that its observed coverage distribution is close to uniform.

At the species level, the simulator recapitulates the qualitative composition of the TELSeq sample. The dominant species in the mock community remains dominant, targeted bacterial species are enriched relative to their theoretical proportions, and non-target fungal and bacterial species remain depleted (Table 4; Supplementary Table S4). The quantitative differences (i.e. underrepresentation of *L. monocytogenes* and overrepresentation of *P. aeruginosa* and *S. enterica* in the simulation) are consistent with real-world experimental biases not incorporated in the current model. In particular, the thermodynamic nearest-neighbor (TNN) implementation used in `RAmpSim` penalizes mismatches by setting their binding energy *ΔG* to 0 rather than mismatch-specific penalties, and GC-related inefficiencies during extraction and library preparation are not modeled. As a result, genomes with higher GC content can be sampled more frequently in silico than they appear in the empirical data, which can suppress the abundance of remaining genomes such as *L. monocytogenes*.

Taken together, these observations point to several extensions. First, incorporating an explicit GC-dependent weighting term, either at the hybridization stage or at a separate amplification stage, should improve the fit for high-GC genomes and reduce peak flattening. Second, replacing the simplified TNN model with a more complete parameterization of sequence-dependent free energies would reduce systematic differences in species-level abundances. These extensions can be added without changing the overall structure of the simulator and while retaining its current runtime characteristics. In practice, this suggests that `RAmpSim` is already well suited for reproducing coarse enrichment structure and qualitative abundance patterns and for benchmarking downstream analyses that depend primarily on relative enrichment.

## 5 Conclusion

In summary, the simulator already reproduces the major spatial and compositional features of the TELSeq dataset while using a single-threaded, memory-resident implementation. This makes it a practical tool for evaluating bait designs, for stress-testing downstream analysis pipelines under controlled conditions, and for studying how individual sources of bias (GC content, bait density, species abundance) propagate to observable coverage. As richer empirical datasets become available, the simulator can be iteratively calibrated using the same framework described here.

## Acknowledgments

A.Z. and Y.W.Y. are supported by the US National Institutes of Health grant R35GM160134. C.B and N.N. are supported by the National Institutes of Health grant R01AI173928. Mock community data were generated under the US National Institutes of Health grant R01AI141810. We also acknowledge Molly Borowiak for fruitful discussions.

