## Supplementary material for "RAmpSim: A Thermodynamic Simulator for Hybridization Capture in Metagenomic Sequencing": Supp_Data

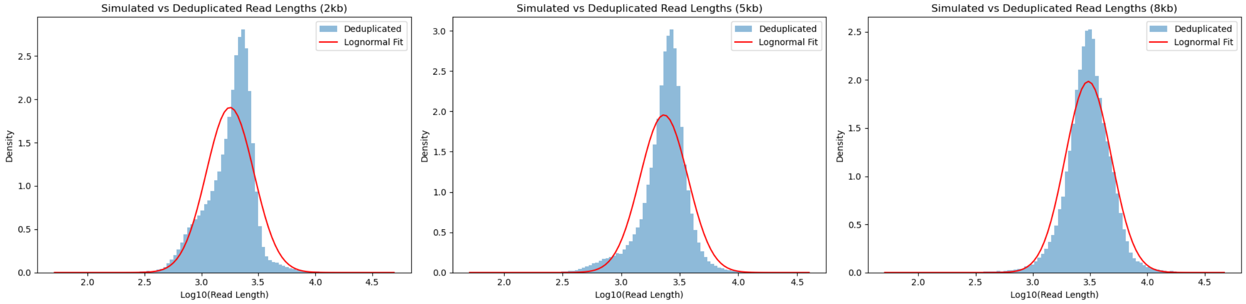


Fig. S1: Fitted lognormal distributions of fragment lengths across replicates. We see that on the log-scale each distribution is approximately normal, and that the model fit improves for larger target fragment lengths. In particular, the model fit is best near the median.

Table S1: Simulation Parameters by Insert Size

| **Insert Size** | **–nfrag** | **–flen** | **–lognorm-sd** | **–split** |
|---|---|---|---|---|
| 8kb | 154000 | 3000 | 0.46 | 0.63 |
| 5kb | 111000 | 2300 | 0.47 | 0.68 |
| 2kb | 125000 | 1800 | 0.48 | 0.85 |

| **Species** | **2kb** | | **5kb baseline** | |
|---|---|---|---|---|
| | **EMD(tels,RAmpSim)** | **EMD(tels,unif)** | **EMD(tels,RAmpSim)** | **EMD(tels,unif)** |
| *Listeria monocytogenes* | **3.30e-07** | 4.98e-07 | **2.33e-07** | 3.20e-07 |
| *Pseudomonas aeruginosa* | **1.21e-07** | 2.68e-07 | **1.55e-07** | 2.75e-07 |
| *Bacillus subtilis* | **5.61e-08** | 4.63e-07 | **8.46e-08** | 4.55e-07 |
| *Escherichia coli* | **2.41e-08** | 3.48e-07 | **2.03e-08** | 3.36e-07 |
| *Enterococcus faecalis* | **1.65e-07** | 6.86e-07 | **2.68e-07** | 6.83e-07 |
| *Lactobacillus fermentum* | **8.58e-07** | 1.05e-06 | **4.49e-07** | 1.05e-06 |
| *Salmonella enterica* | **7.83e-08** | 3.63e-07 | **7.25e-08** | 3.55e-07 |
| *Staphylococcus aureus* | **5.51e-07** | 7.33e-07 | **6.12e-07** | 7.34e-07 |

Table S2: EMD results for the 2 kb and 5kb replicates. For prokaryotic genomes with single chromosomes, the EMD between the empirical and simulated coverage distributions outperforms the baseline. Eukaryotic species such as *S. cerevisiae* and *C. neoformans* contain multiple chromosomes of varying lengths and were not considered targets in the original bait design.

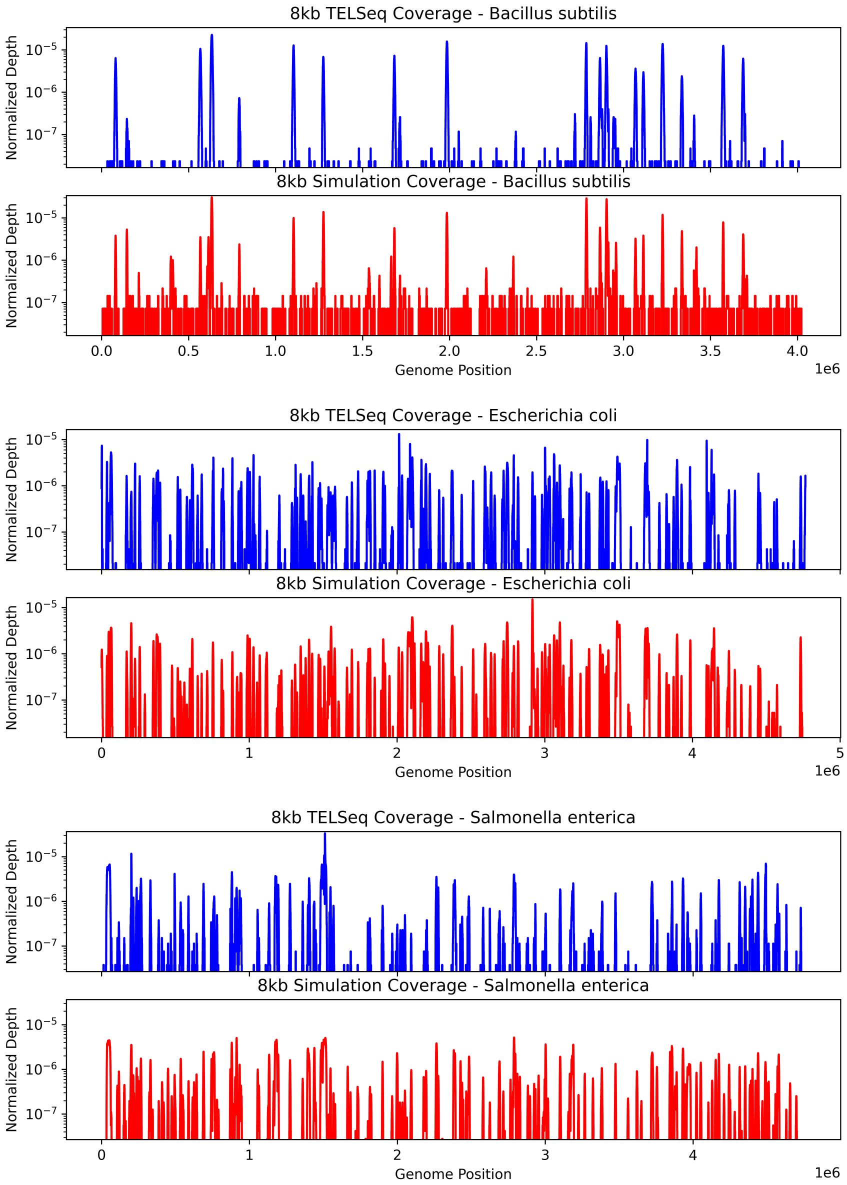


Fig. S2: Log coverage distribution plots for *B. subtilis* (top), *E. Coli* (middle), and *S. enterica* (bottom). The coverage distribution for each genome was normalized by the sum of the coverage over each position. The coverage patterns between the observed (blue) and simulated (red) data are similar, in which peaks in each distribution correspond to high bait-density regions shown in gray. Outside of these regions, coverage falls down to background levels.

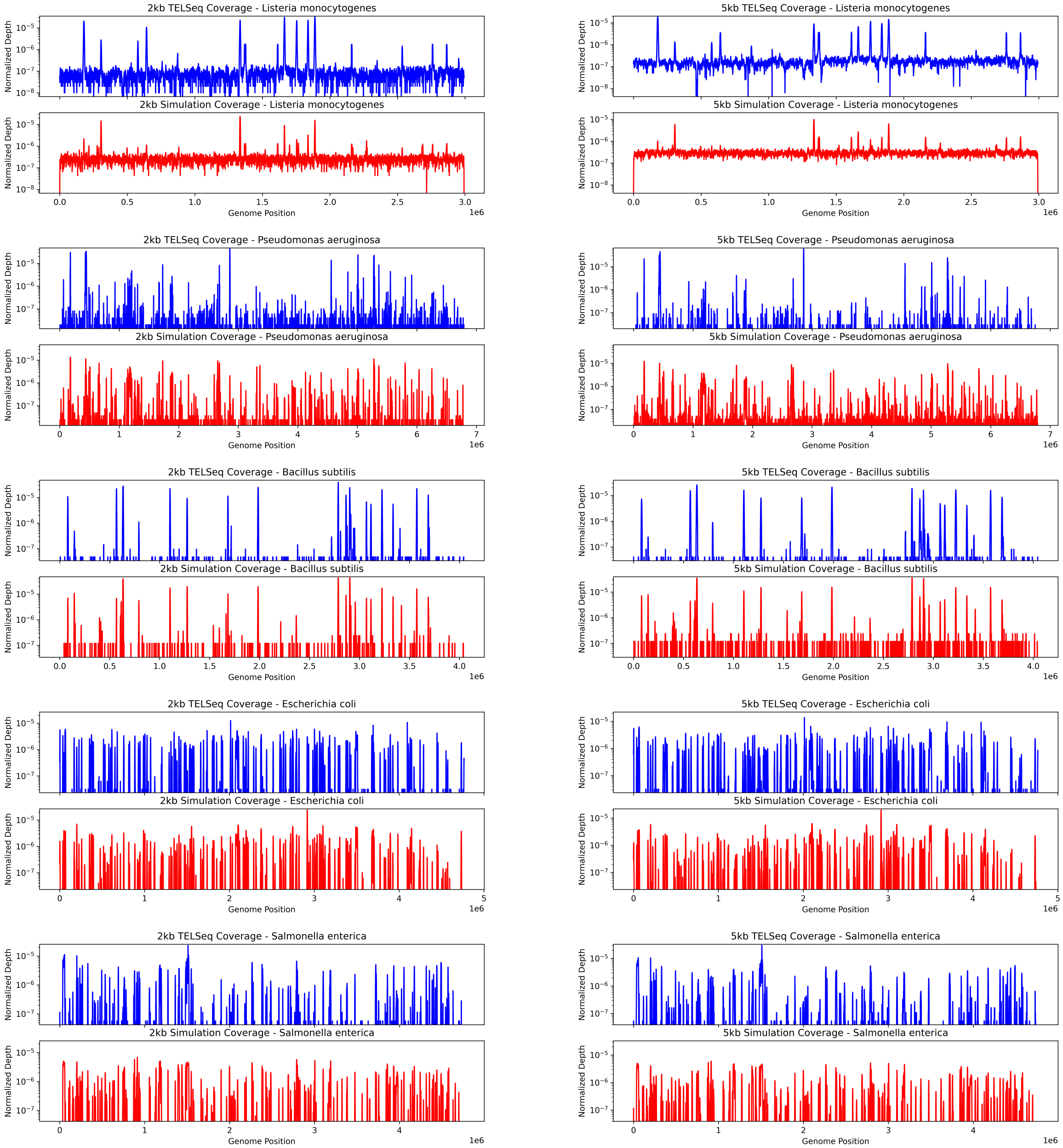


Fig. S3: Log coverage distribution plots of large genomes for 2kb and 5kb replicates. The coverage distribution for each genome was normalized by the sum of the coverage over each position. Similar to the coverage patterns in the 8kb replicates, the observed (blue) and simulated (red) data for the 2kb and 5kb replicates share similar coverage patterns.

| Species | RAmpSim (2kb) | | | Uniform baseline (2kb) | | |
|---|---|---|---|---|---|---|
| | **F1 Score** | **Precision** | **Recall** | **F1 Score** | **Precision** | **Recall** |
| *Listeria monocytogenes* | **0.4864** | **0.5023** | **0.4714** | $0.453 \pm 0.022$ | $0.461 \pm 0.028$ | $0.446 \pm 0.019$ |
| *Pseudomonas aeruginosa* | 0.3269 | 0.3156 | **0.3391** | $0.318 \pm 0.014$ | $0.447 \pm 0.022$ | $0.247 \pm 0.011$ |
| *Bacillus subtilis* | **0.3570** | 0.3663 | **0.3480** | $0.196 \pm 0.024$ | $0.405 \pm 0.052$ | $0.129 \pm 0.015$ |
| *Escherichia coli* | **0.7785** | **0.8296** | **0.7333** | $0.289 \pm 0.035$ | $0.409 \pm 0.095$ | $0.225 \pm 0.018$ |
| *Salmonella enterica* | **0.7286** | **0.8710** | **0.6263** | $0.245 \pm 0.034$ | $0.363 \pm 0.115$ | $0.187 \pm 0.016$ |
| *Lactobacillus fermentum* | 0.0000 | 0.0000 | 0.0000 | $0.000 \pm 0.000$ | $0.000 \pm 0.000$ | $0.000 \pm 0.000$ |
| *Enterococcus faecalis* | **0.4742** | **0.9999** | **0.3108** | $0.036 \pm 0.040$ | $0.087 \pm 0.097$ | $0.023 \pm 0.026$ |
| *Staphylococcus aureus* | 0.1199 | **0.9641** | 0.0639 | $0.059 \pm 0.086$ | $0.060 \pm 0.084$ | $0.059 \pm 0.088$ |

| Species | RAmpSim (5kb) | | | Uniform baseline (5kb) | | |
|---|---|---|---|---|---|---|
| | **F1 Score** | **Precision** | **Recall** | **F1 Score** | **Precision** | **Recall** |
| *Listeria monocytogenes* | 0.5033 | 0.5013 | 0.5052 | $0.477 \pm 0.027$ | $0.467 \pm 0.039$ | $0.488 \pm 0.019$ |
| *Pseudomonas aeruginosa* | **0.2576** | **0.5578** | **0.1675** | $0.200 \pm 0.017$ | $0.379 \pm 0.033$ | $0.136 \pm 0.012$ |
| *Bacillus subtilis* | **0.3575** | 0.4528 | **0.2953** | $0.230 \pm 0.024$ | $0.412 \pm 0.046$ | $0.160 \pm 0.016$ |
| *Escherichia coli* | **0.7731** | **0.8023** | **0.7459** | $0.344 \pm 0.022$ | $0.479 \pm 0.032$ | $0.268 \pm 0.018$ |
| *Salmonella enterica* | **0.7140** | **0.8541** | **0.6134** | $0.267 \pm 0.028$ | $0.353 \pm 0.038$ | $0.215 \pm 0.022$ |
| *Lactobacillus fermentum* | 0.0162 | 0.0682 | 0.0092 | $0.008 \pm 0.068$ | $0.007 \pm 0.064$ | $0.009 \pm 0.072$ |
| *Enterococcus faecalis* | **0.5064** | **0.9997** | **0.3391** | $0.051 \pm 0.028$ | $0.246 \pm 0.133$ | $0.028 \pm 0.016$ |
| *Staphylococcus aureus* | 0.0805 | **0.9702** | 0.0420 | $0.070 \pm 0.108$ | $0.067 \pm 0.105$ | $0.075 \pm 0.111$ |

Table S3: F1, precision, and recall scores of high/low coverage peak classification by RAmpSim and a uniform baseline classifier in the 2kb replicate (top) and 5kb replicate (bottom). Uniform baseline values are reported as mean ± CI. Compared to the 8kb replicate (Table 3), a larger number of species in the 2kb and 5kb replicates have RAmpSim F1, precision, and recall values that are close to, or fall within, the confidence intervals of the uniform baseline. This includes *P. aeruginosa* and *B. subtilis*, which exhibit more prominent background-like coverage distributions in the observed data. Performance similarly degrades for extremely low-abundance species such as *S. aureus*, where RAmpSim primarily struggles with recall, and for untargeted species such as *L. fermentum*, which show very little enrichment signal above background.

Table S4: Comparison of Theoretical, Observed, and Simulated Values for the 2kb and 5kb Replicates.

| **Species** | **Theoretical** | **2kb** | | **5kb** | |
|---|---|---|---|---|---|
| | | **Obs** | **RAmpSim** | **Obs** | **RAmpSim** |
| *Listeria monocytogenes* | 89.100000 | 44.255939 | 17.737358 | 57.308454 | 31.895452 |
| *Pseudomonas aeruginosa* | 8.900000 | 22.881611 | 33.129358 | 13.044702 | 28.086749 |
| *Bacillus subtilis* | 0.890000 | 9.634952 | 3.345772 | 9.719870 | 2.936187 |
| *Saccharomyces cerevisiae* | 0.890000 | 0.085608 | 0.205342 | 0.098807 | 0.304543 |
| *Escherichia coli* | 0.089000 | 14.470449 | 19.634965 | 12.186773 | 15.969443 |
| *Salmonella enterica* | 0.089000 | 8.635885 | 20.150204 | 7.609556 | 16.146896 |
| *Lactobacillus fermentum* | 0.008900 | 0.000564 | 0.204294 | 0.000266 | 0.185552 |
| *Enterococcus faecalis* | 0.000890 | 0.024360 | 2.340124 | 0.023562 | 1.844750 |
| *Cryptococcus neoformans* | 0.000890 | 0.000000 | 0.000000 | 0.000000 | 0.000000 |
| *Staphylococcus aureus* | 0.000089 | 0.010067 | 3.048291 | 0.007745 | 2.444875 |
